# Viral long-term evolutionary strategies favor stability over proliferation

**DOI:** 10.1101/539239

**Authors:** Stéphane Aris-Brosou, Louis Parent, Neke Ibeh

## Abstract

Viruses are known to have some of the highest and most diverse mutation rates found in any biological replicator, with single-stranded (ss) RNA viruses evolving the fastest, and double-stranded (ds) DNA viruses having rates approaching those of bacteria. As mutation rates are tightly and negatively correlated with genome size, selection is a clear driver of viral evolution. However, the role of intragenomic interactions as drivers of viral evolution is still unclear. To understand how these two processes affect the long-term evolution of viruses infecting humans, we comprehensively analyzed ssRNA, ssDNA, dsRNA, and dsDNA viruses, to find which virus types and which functions show evidence for episodic diversifying selection and correlated evolution. We show that selection mostly affects single stranded viruses, that correlated evolution is more prevalent in DNA viruses, and that both processes, taken independently, mostly affect viral replication. However, the genes that are jointly affected by both processes are involved in key aspects of their life cycle, favoring viral stability over proliferation. We further show that both evolutionary processes are intimately linked at the amino acid level, which suggests that it is the joint action of selection and correlated evolution, and not just selection, that shapes the evolutionary trajectories of viruses – and possibly of their epidemiological potential.

## Introduction

Humanity is regularly reminded of the epidemiological toll of viruses, in part due to recent and ongoing viral outbreaks of influenza Smith *et al.* (2009), Ebola Gire *et al.* (2014), and Zika Faria *et al.* (2016a). Thanks to recent technological and analytical developments, it is now possible to elucidate, almost in real time Faria *et al.* (2016b), their epidemiological dynamics, and unravel their evolutionary dynamics Grenfell *et al.* (2004). However, while the evolutionary dynamics of RNA viruses are well documented Holmes (2009), those of other viruses are not so well known, in particular in a unifying context including major viral types such as double-stranded (ds) and single-stranded (ss) DNA and RNA viruses.

To date, one of the most salient evolutionary feature shared by all viruses is the existence of a negative correlation between mutation rate and genome size Holmes (2009). This is a critical result as it suggests that selection is driving the evolution of mutation rates, establishing a tradeoff between mutational load and availability of adaptive mutations Holmes (2011). But the processes driving the evolution of different types of viruses are multiple. As already argued, both ssDNA and ssRNA viruses share small genome sizes, high mutation rates, but also large effective population sizes, little to no gene duplication or recombination, and overlapping reading frames Holmes (2009). This last point suggests that a less frequently explored evolutionary process in viral studies, correlated evolution, could be as critical as positive selection. Correlated evolution happens when mutations at two locations in a genome occur one after the other, in a quick succession Kryazhimskiy *et al.* (2011), repeatedly Maddison and FitzJohn (2015). Typical examples include drug resistance mutations that have a fitness cost, and that require a second mutation to compensate for the first one Gong *et al.* (2013), or tRNAs that require a specific base-pairing to maintain their secondary and tertiary structures, so that a mutation in the stem region necessitates a second mutation to restore the correct, functional, structure Meer *et al.* (2010). This process is of particular interest as correlated evolution can be underlain by epistasis, which occurs when the fitness effects of these two mutations are non-additive Dench *et al.* (2019), as in the two examples above. As such, correlated evolution can help us understand the relationship between genotype and fitness, which is a key determinant of evolutionary trajectories Li *et al.* (2016). To date however, correlated evolution has only sporadically been investigated in viral evolution, and these rare instances only focused on ssRNA viruses. Indeed, recent work uncovered pervasive evidence for correlated evolution in influenza viruses Nshogozabahizi *et al.* (2017); Lyons and Lauring (2018), and both the Zika Aris-Brosou *et al.* (2017) and the Ebola viruses Ibeh *et al.* (2016). Intriguingly, in this latter case (Ebola), evidence was found that sites evolving in a correlated manner could also be under positive selection – bearing the question as to how frequently these two processes, correlated evolution and positive selection, occur, possibly jointly, and if this co-occurrence is limited to ssRNA viruses, or can be generalized to all viruses.

To better understand the role of correlated evolution and positive selection in the evolutionary dynamics of viruses infecting humans, we constructed a nearly exhaustive viral data set spanning all dsDNA, dsRNA, ssRNA, and ssDNA viruses deposited in GenBank (as of August 2017), and conducted an extensive survey of correlated evolution and diversifying selection in these viruses, asking more specifically about the prevalence of these two processes in each viral type, independently or jointly, with the specific hypothesis that the genes affected by both processes encode functions that are most critical to each viral life cycle.

## 1 Materials and Methods

### 1.1 Data retrieval

Lists of dsDNA, dsRNA, ssRNA, and ssDNA viruses infecting humans were retrieved from the viruSITE database Stano *et al.* (2016) in August 2017 (Tables S1-S4); subtypes / genotypes / clades were treated as independent data sets. Although some of these viruses could be segmented or not, with circular or linear genome, with positive or negative strands, and with or without overlapping reading frames, accounting for these structural features would have led to smaller and smaller data sets, precluding any statistical analysis, so that the data were not split beyond viral type. Each list contained the virus names, the length of their genome, their number of protein-coding genes (CDS’s), and was associated with a reference coding sequence (see `query sequences.zip` at https://github.com/sarisbro). In order to obtain corresponding sequence alignments of orthologous genes, BLASTn searches were performed on a custom database limited to viral genes present in NCBI nucleotide database with blast-2.6.0+. For this, all gbvrl*.seq.gz files were downloaded from ftp.ncbi.nih.gov/genbank while querying viruSITE, and were concatenated into a single GenBank file, then converted into a FASTA file with `readseq` to specifically extract CDS’s Gilbert (2003). This was done to avoid retrieving 5’ and 3’ untranslated regions that would cause problems to the downstream codon analyses. BLASTn searches were performed for each viruSITE viral sequence with a stringent E-value threshold of 10^−100^, keeping a maximum of 100 sequences with at least 80% coverage with each query; this ensured that subtype / genotype / clade boundaries were not crossed. As the viruses retrieved from the viruSITE also included viruses that require a vector (*e.g.*, arboviruses such as the Dengue and Yellow fever viruses), or viruses that circulate in non-human hosts but that can lead to zoonoses (*e.g.*, the Camel alphacoronavirus leading to MERS van Boheemen *et al.* (2012); Zaki *et al.* (2012)), these stringent thresholds also ensured that sequences contained in each alignment mostly came from a single host. Only data sets with at least 20 hits were kept for downstream analyses. Sequences corresponding to each accession number were retrieved from the FASTA file obtained with `readseq` Gilbert (2003).

Because this file contained partial sequences, each set of retrieved sequences was first aligned with `Muscle` 3.8.31 Edgar (2004). Each alignment was then quality checked ensuring that (i) its length is a multiple of three, (ii) it starts with an ATG and stops with a stop codon. Alignments failing at least one condition were discarded. Within each alignment, mean numbers of indels 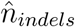 were computed for each sequence, and those containing number of indels 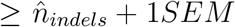 (standard error of the mean) were eliminated. The remaining nucleotide sequences were then re-aligned with TranslatorX Abascal *et al.* (2010), at the amino acid level, using the `Muscle` aligner and their heuristics to determine the correct reading frame. Alignment files were then cleaned-up with `Gblocks` 0.91b at the codon level using the stringent default settings Castresana (2000). Both trees and alignments are available from https://github.com/sarisbro.

To obtain gene annotations, Gene Ontology (GO) terms were retrieved from gene sequences with HMMER2GO (github.com/sestaton/HMMER2GO), relying on the Hidden Markov Models in ftp.ebi.ac.uk/pub/databases/Pfam/current_release/Pfam-A.hmm.gz (ver. Feb 23, 2017). This is equivalent to performing a Blast2GO search Conesa *et al.* (2005), but without the limitations of proprietary software. Individual mapping files coming from our four viral types (ssRNA, dsRNA, ssDNA, and dsDNA viruses) were then merged to create a custom annotation file, used for GO term enrichment testing with topGO Alexa and Rahnenführer (2009), based on Fisher’s exact test. The reference gene list was always the entire set of genes within each viral type.

### 1.2 Phylogenetic analyses

Phylogenetic trees were reconstructed with `FastTree` 2.1.7 Price *et al.* (2010) under the GTR+Γ model of evolution Aris-Brosou and Rodrigue (2012); note that `FastTree` was recompiled locally to use double-precision arithmetics, as recommended by its authors to estimate very short branch lengths accurately. Those with a nonzero tree length were midpoint rerooted using the phytools package ver. 0.4-60 Revell (2012) in R ver. 3.2.3 R Core Team (2016).

Patterns of correlated evolution among sites were identified with the Bayesian graphical model (BGM) implemented in SpiderMonkey Poon *et al.* (2008), which is part of HyPhy ver. 2.3.3 Pond *et al.* (2005). Default HyPhy scripts were slightly modified to read in codon data, which were used to reconstruct mutational paths under the MG94×HKY85 substitution model Kosakovsky Pond and Frost (2005) at nonsynonymous sites along each branch of the estimated trees. Ambiguous reconstructions were resolved by considering all possible resolutions and averaging them. These reconstructed mutational paths were then recoded as a binary matrix, with rows corresponding to branches and columns to a site of the alignment. The BGM was then used to identify the pairs of sites that exhibit correlated patterns of nonsynonymous substitutions according to their posterior probability, estimated with a Markov chain Monte Carlo sampler that was run for 10^5^ steps, with a burn-in period of 10,000 steps sampling every 1,000 steps for inference Aris-Brosou *et al.* (2017).

Patterns of episodic selection were identified based on the Mixed Effects Model of Evolution Murrell *et al.* (2012), also as implemented in HyPhy. The default script from ver. 2.2.6 was used, still with the 2.3.3 HyPhy engine, to infer nonsynonymous to synonymous rate ratios *ω* assuming that these rates can vary across lineages and among sites. In this implementation, two categories of sites were assumed, those for which *ω*_*neg*_ ≤ 1, in proportion *p*, and those for which *ω_pos_* > 1, that are under positive selection, in proportion 1 − *p*. Evidence for selection was derived by means of a likelihood ratio test between this model, and a null model where *ω_pos_* was constrained to take its value between 0 and 1. Linear models were fitted through robust regressions Yohai *et al.* (1991). All R scripts and HyPhy source files are available from https://github.com/sarisbro (file “API_scripts.zip,” in “data”).

## 2 Results and Discussion

First, in order to better understand the genomic characteristics of the four types of viruses known to infect humans, dsDNA (*n* = 94), dsRNA (*n* = 15), ssRNA (*n* = 354), and ssDNA (*n* = 84), and understand how these characteristics can impact the evolutionary dynamics of these viruses (Methods; Figure S1), we examined the distribution of four of their genomic features. While dsDNA viruses had the largest and the most variable genome lengths, ssDNA were the smallest genomes by at least an order of magnitude, with both dsRNA and ssRNA exhibiting intermediate sizes (Figure 1a). Because of the compact structure of these genomes, these differences were also reflected in the number of genes, protein-coding genes, and RNAs encoded by the genomes of these viruses (Figure S2). In turn, these characteristics suggest that with their large genomes, dsDNA viruses potentially have a higher degree of functional redundancy than ssDNA or even ssRNA viruses, possibly due to duplication events Gao *et al.* (2017), and thereby be under less stringent selective pressures than viruses with smaller genomes. On the other hand, the extent of correlated evolution and its interaction with selection is more difficult to predict.

**Figure 1.**
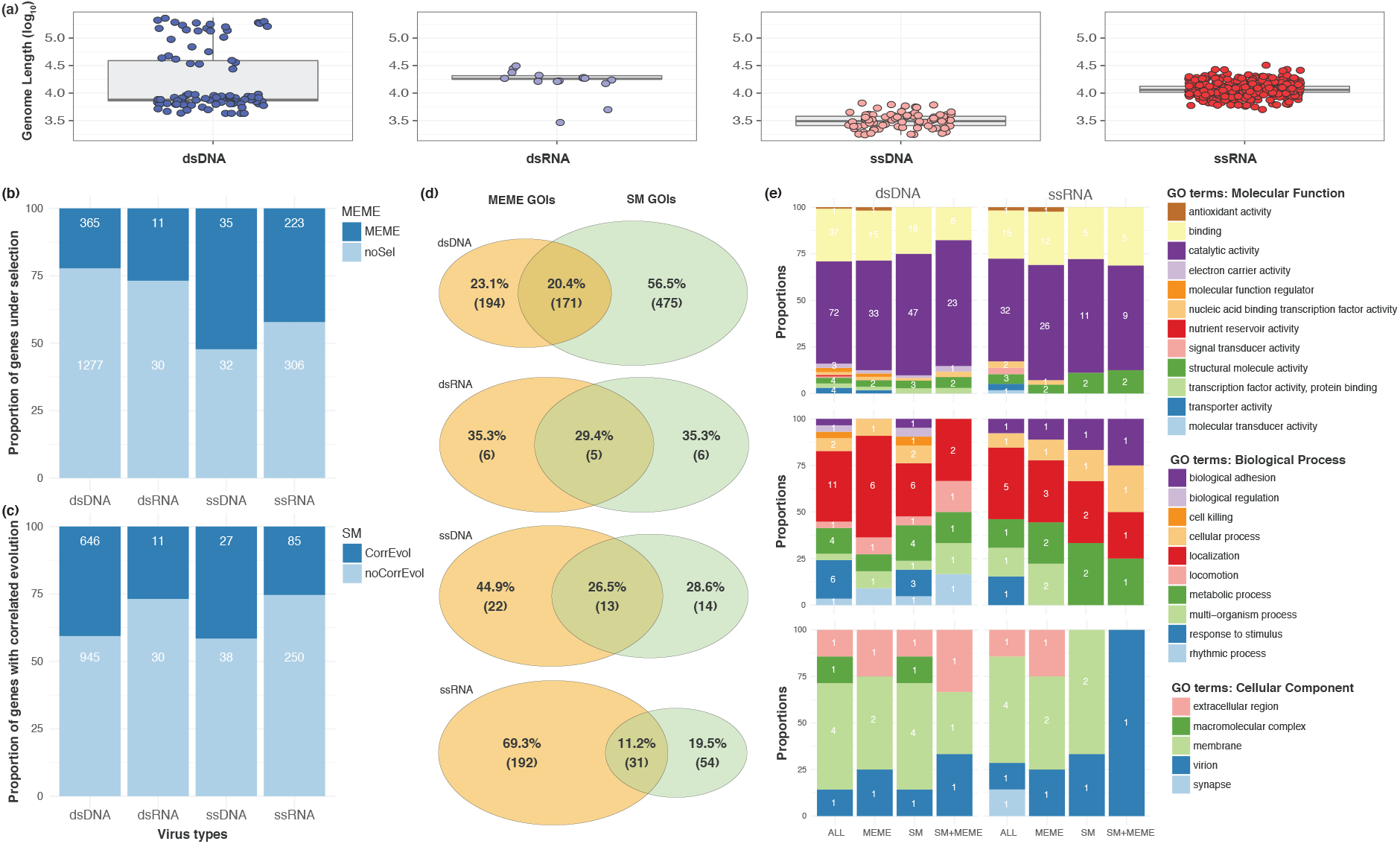
Modes of evolution of viruses. Modes of evolution of viruses. (**a**) Distribution of genome size, on a log_10_ scale ssRNA (blue), ssDNA (purple), dsRNA (orange), and dsDNA (red). (**b**) Proportion of genes detected to be under diversifying selection (top, dark hues); actual numbers of genes are shown within each column. (**c**) Proportion of genes detected to be evolving in a correlated manner (top, dark hues); actual numbers of genes are shown within each column. (**d**) Venn diagrams showing the Genes of Interest (GOIs) either under selection (MEME GOIs), or evolving in a correlated manner (SM GOIs), or both (intersect) for each virus type. (**e**) Gene Ontology (GO) enrichment tests for the genes that are both under selection and evolving in a correlated manner for their Molecular Function (top), Biological Processes (middle), and Cellular Component (bottom), all at level 2 of the ontology.

To address these questions, we first tested for the presence of episodic diversifying selection in each viral type. Altogether, we found extensive differences among all four viral types in terms of the number of genes under selection (Figure 1b; *X*^2^ = 99.61, *df* = 3, *P* < 2.2 × 10^−16^), and that single stranded viruses were more subject to selection than double stranded viruses (*X*^2^ = 95.14, *df* = 1, *P* < 2.2 × 10^−16^). Indeed, and *contra* our original hypothesis, there were no differences between dsDNA and dsRNA viruses (*X*^2^ = 0.26, *df* = 1, *P* = 0.6110), or between ssDNA and ssRNA viruses in terms of prevalence of diversifying selection (*X*^2^ = 2.07, *df* = 1, *P* = 0.1503). Note that these differences cannot be attributed to genetic diversity, as dsDNA and dsRNA, which have similar levels of selection, have however different levels of diversity (Figure S3). However, it is unlikely that “strandedness” (single *vs.* double stranded genetic material) alone drives selection, even if greater instability can be postulated in single-stranded nucleic acids Frederico *et al.* (1990). Indeed, strandedness is negatively correlated with genome size (*t* = −4.98, *df* = 108.35, *P* = 2.43 × 10^−6^), which is itself correlated with mutation rates Lynch (2007); Duffy (2018). As a result, episodic diversifying selection is mostly driven by structural aspects of viral genomes, which condition mutation rates across all virus types Sanjuán (2012).

To understand if similar aspects drive intragenic correlated evolution, we counted in the same way the number of genes for which we could find evidence for interactions. Here again, we found extensive differences among all four viral types (Figure 1c; *X*^2^ = 29.79, *df* = 3, *P* < 1.5 × 10^−16^), with no difference between dsDNA and ssDNA viruses (*X*^2^ = 0.0005, *df* = 1, *P* = 0.9827), or between dsRNA and ssRNA viruses (*X*^2^ = 0.0001, *df* = 3, *P* = 0.9903). Again, diversity is not driving these differences, as both dsDNA / ssDNA and dsRNA / ssRNA have significantly different levels of diversity (Figure S3). As a result, the prevalence of correlated evolution seems to be mostly driven by the nature of viral genetic material. At least two processes can underpin correlated evolution: linkage and epistasis Dench *et al.* (2019). As recombination is pervasive in some dsDNA viruses Robinson *et al.* (2011), linkage alone may not explain the high prevalence of correlated evolution in DNA viruses (close to 40%: Figure 1c). Rather, this pattern suggests that intragenic constraints are higher in DNA viruses for unknown structural reasons, maybe due to protein structure Woo *et al.* (2014), or in the same way that recombination in RNA viruses may be driven by mechanistic constraints associated with genome structures and viral life cycles Simon-Loriere and Holmes (2011).

While previous work showed that both diversifying selection and correlated evolution can affect the same gene and even the same site in a viral genome such as Ebola’s Ibeh *et al.* (2016), a ssRNA virus, the generality of this association is still unknown. At the gene level, we found strong heterogeneity among all four viral types for gene numbers showing evidence for selection, correlated evolution or both (*X*^2^ = 206.54, *df* = 6, *P* < 2.2×10^−16^). Surprisingly, ssRNA viruses are those that show the least overlap between selection and correlated evolution, with only 11% of the genes evolving under both mechanisms (Figure 1d). Patterns are however much more difficult to extract here, mostly because the number of genes involved becomes quite small, in particular for the virus type with most overlap, the dsRNA viruses (Figure 1d: 29.4%, *i.e.* five genes).

To assess the extent to which some of these differences at the gene level are functionally driven, we extracted the Gene Ontology (GO) annotations, or GO terms, associated with the genes analyzed, as well as those under selection, correlated evolution, or both – focusing exclusively on the virus types for which we had the largest samples sizes, dsDNA and ssRNA viruses (Figure 1e). Each of the three parts of the ontology (Molecular Function [MF], Biological Process [BP], and Cellular Component [CC]) were first limited to the second level of GO in order to derive a high-level interpretation (low-level descriptions are shown in Tables S1-S3). Figure 1e shows that GO terms related to catalytic activity (MF), involved in the establishment or maintenance of a certain location (BP) at the membrane level (CC) are predominantly present in both dsDNA and ssRNA genomes. However, only a subset of these GO terms is mostly under episodic diversifying selection: helicase activity / binding (MF) and replication (BP) in the host cell nucleus (CC) in dsDNA, while it is mostly peptidase activity and methylation on the viral envelope in ssRNA viruses (Table S1; *P* < 0.01). In spite of these differences, we note that episodic diversifying selection mostly affects genes involved in viral replication (Table S1). Genes that are affected by correlated evolution include: helicase activity and binding at the interface of multiple compartments in dsDNA viruses, or transferase activity and protein modifications on the envelope in ssRNA viruses (Table S2). Again, most of these functions and processes are involved in viral replication (Table S2). At the intersect of these evolutionary processes however, the genes that are jointly affected by selection and correlated evolution are involved in structural integrity and assembly within or outside a cell at the capsid level for dsDNA viruses, or in interacting with host cell surface via antigen activity on the viral envelope for ssRNA viruses – functions and processes that are mostly involved in “viral stability” (cell entry, integrity, assembly, immune escape; Table S3). This suggests that despite key differences in life history strategies adopted by dsDNA and ssRNA, there is a certain unity across viral types, where genes involved in replication are mostly under either selection or correlated evolution, while those involved in viral stability are mostly affected by both evolutionary processes – as has been shown for the influenza Gong *et al.* (2013) and the Ebola Ibeh *et al.* (2016) viruses (both cases involved a glycoprotein required for cell entry). Because these two evolutionary processes are required by epistasis, it is possible that the genes involved in viral stability are the most likely to show evidence for non-additive fitness effects – and hence, shape viral fitness landscapes. This tension between replication and stability is evocative of the existence of tradeoffs between capsid stability and proliferation within a host De Paepe and Taddei (2006), or fecundity and lifespan García-Villada and Drake (2013) in dsDNA viruses, which supports the idea that it is the joint action of selection and of correlated evolution that shapes viral life histories.

The previous analyses were at the gene level. One outstanding question is whether this relationship between selection and correlated evolution also holds at the amino acid level, that is, if the amino acids under selection are also involved in correlated evolution. This question was addressed in two different ways, all virus types confounded (in order to have larger sample sizes), by focusing on one process at a time, and finding at what point evidence supporting the second process becomes significant. First, we searched the list of pairs of sites evolving in a correlated manner (the SM sites) to see if at least one pair member was potentially evolving under episodic diversifying selection (the MEME sites), irrespective of its probability of being a MEME site. For this, we identified pairs of SM sites, that is those with a posterior probability ≥ 0.95, and plotted their posterior probability as a function of the − log_10_ probability of each pair member to be under selection (Figure 2a). Both the least-square (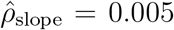, *t* = 3.90, *P* = 0.0001) and the robust regressions (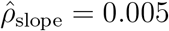, *t* = 2.62, *P* = 0.0089) have positive and significant slopes, hereby demonstrating the existence of a relationship between correlated evolution and episodic diversifying selection at the amino acid level – even if most of the SM sites show weak evidence of selection (at *P* ≤ 0.01, gray broken line in Figure 2a). So, to validate the existence of this relationship, we then took the list of MEME sites, and searched the list of SM sites to see if at least one pair member was potentially a MEME site, irrespective of its posterior probability of being in an SM site pair. For this, we identified the MEME sites, that is those with a *P*-value ≤ 0.01 (≥ 2 on a − log_10_ scale), and plotted this (on the *x*-scale) *vs.* their posterior probability of being in an SM pair (on the *y*-scale; Figure 2b). Both the least-square (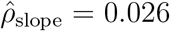, *t* = 3.64, *P* = 0.0003) and the robust regressions (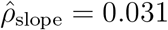, *t* = 3.37, *P* = 0.0008) have positive and significant slopes, further confirming the existence of a relationship between correlated evolution and episodic diversifying selection at the amino acid site level. Note however that this latter analysis is more informative than the former, as the density also shows that most of the sites under selection are involved in weak interactions. This is intriguingly reminiscent of the involvement of weakly interacting pairs of sites in severe outbreaks or pandemics Aris-Brosou *et al.* (2017).

**Figure 2.**
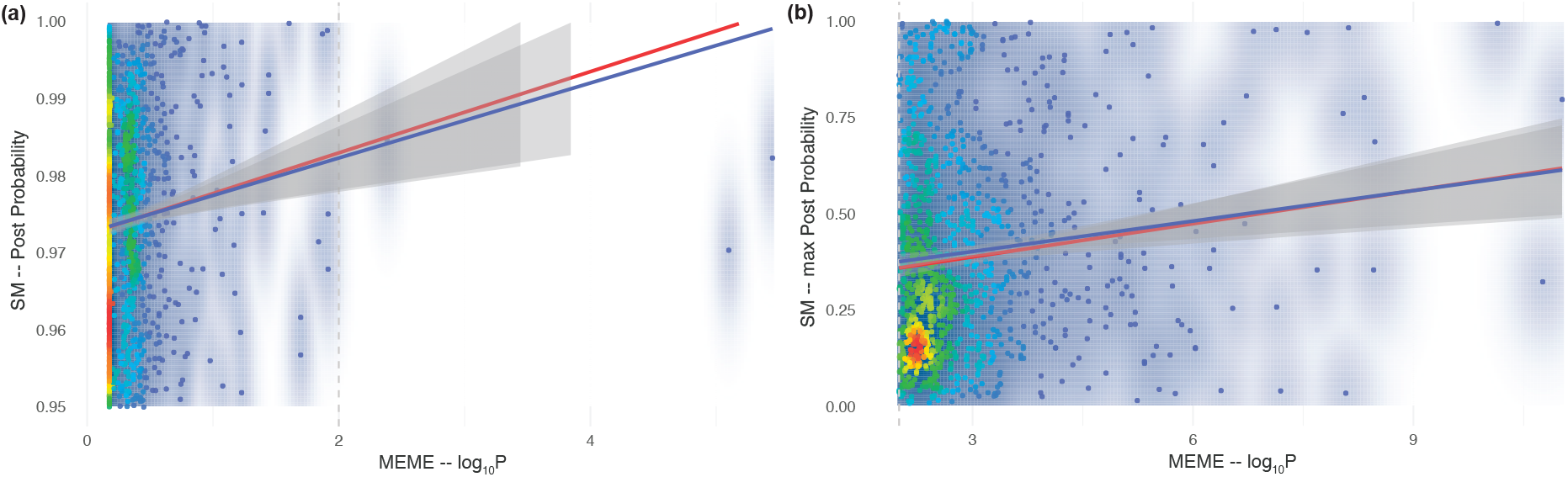
Strength of evidence for sites under both selection (MEME) and correlated evolution (SM). (**a**) SM sites that are also under episodic diversifying selection. (**b**) MEME sites that are also evolving in a correlated manner. In both cases, the posterior probability of a site evolving in a correlated manner is plotted as a function of the *P*-value of a site being under selection (on a − log_10_ scale). Regular least-square regressions are shown in blue; robust regressions are shown in red; 95% confidence envelopes are shaded in grey. See text for statistical details. Density scale: from low (cool colors) to high (warm colors).

## 3 Conclusions

Altogether, we showed that episodic diversifying selection is mostly found in single stranded viruses, while correlated evolution is more prevalent in DNA viruses. More critically, we also showed that the genes affected by each process, when acting independently, are involved in viral replication. However, the genes that are jointly affected by both processes are mostly involved in viral stability (cell entry, integrity, assembly, immune escape), and that the same amino acid sites tend to be affected by both processes. In retrospect, this tight relationship between selection and correlated evolution may not be surprising, as correlated evolution can be underlain by epistasis Nshogozabahizi *et al.* (2017); Dench *et al.* (2019), which directly involves selection (in a non-additive way). Epistasis being a key determinant of fitness landscapes and hence, of evolutionary trajectories Li *et al.* (2016), our results suggest that in the long-term, both processes jointly shape the life history of viruses, favoring stability over proliferation. If so, analyzing viral evolution in the joint light of selection and correlated evolution might help us better predict how viruses that affect humans might evolve Weinreich *et al.* (2006); Pedruzzi *et al.* (2018) – as predicting their evolution Sandie and Aris-Brosou (2014) and epidemiology Ben-Nun *et al.* (2019) has a long history fraught with mixed success.

We note however that we neglected some aspects of viral structure: indeed, viruses can be segmented or not, with a circular or linear genome, with positive or negative strands, overlapping reading frames, complications that we could not consider here due to the resulting small sample sizes, even if these factors can impact the mode of evolution of viruses Holmes (2009). Future work should however strive to address these limitations. We also neglected the population genetics context in which different viruses evolve, context that can often be correlated to structural constraints Lynch (2007); Holmes (2009). Furthermore, as we solely focused on intragenic interactions, and not intergenic or higher order correlations, it is not impossible that we missed higher-level constraints affecting viral evolution. In particular, it is possible that intergenic correlations reveal the nature of tradeoffs shaping the life history strategies in viruses De Paepe and Taddei (2006). Future modeling Neverov *et al.* (2015) and empirical Ashenberg *et al.* (2017) work should probably focus on elucidating not only these constraints, but also the theoretical basis connecting correlated evolution, if not epistasis, to selection in shaping the evolutionary strategies of biological replicators, as current evidence linking these processes is currently limited to the influenza Gong *et al.* (2013) and the Ebola Ibeh *et al.* (2016) viruses.

## 4 Acknowledgements

We thank Compute Canada and Ontario’s Centre for Advanced Computing for giving us access to their servers. This work was supported by the Natural Sciences Research Council of Canada (SAB), and by the University of Ottawa (LP, NI).

