## Supplementary Material for "Viral long-term evolutionary strategies favor stability over proliferation"

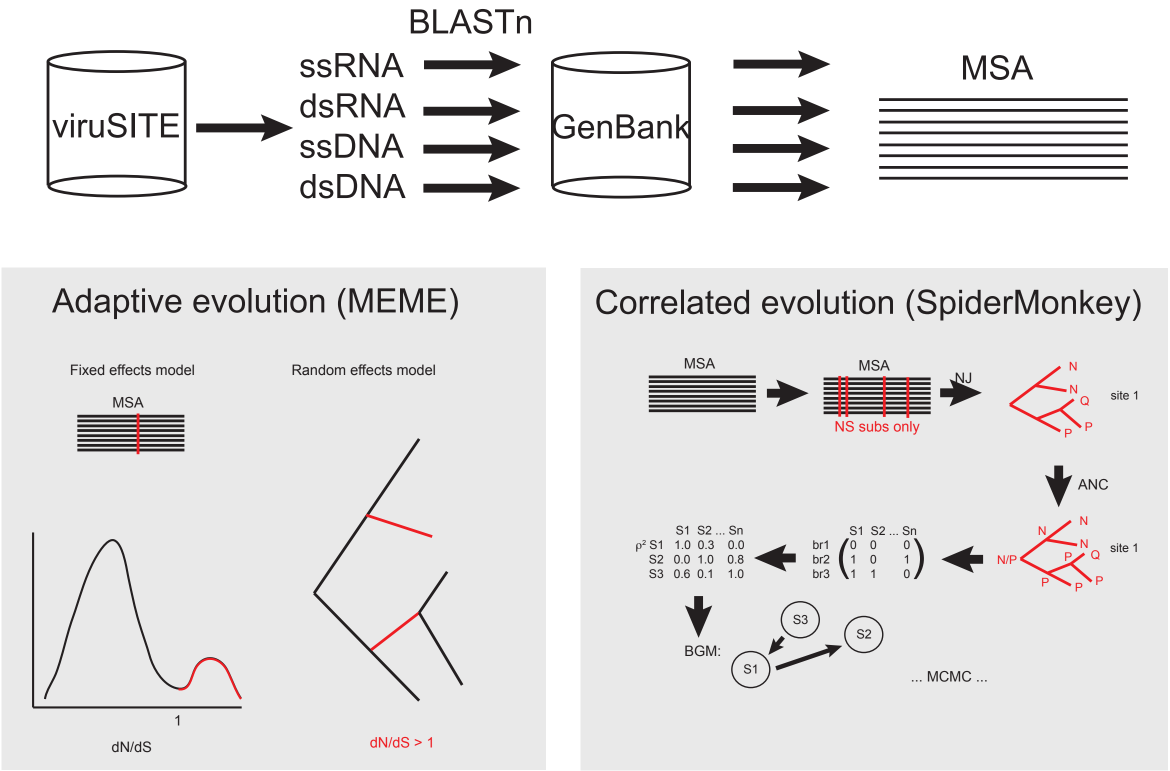


**Figure S1: Overview of the experimental procedures.** Nucleotide sequences of the four types of viruses (dsDNA, dsRNA, ssRNA, and ssDNA) were extracted from viruSITE, and stringent BLASTn searches were conducted on a custom GenBank NR database. These sequences were then aligned, and these multiple sequence alignments (MSA) were subjected to two independent sets of analyses: one searching for evidence for adaptive (diversifying) selection based on the MEME algorithm, and another one searching for evidence for correlated evolution based on the SpiderMonkey algorithm.

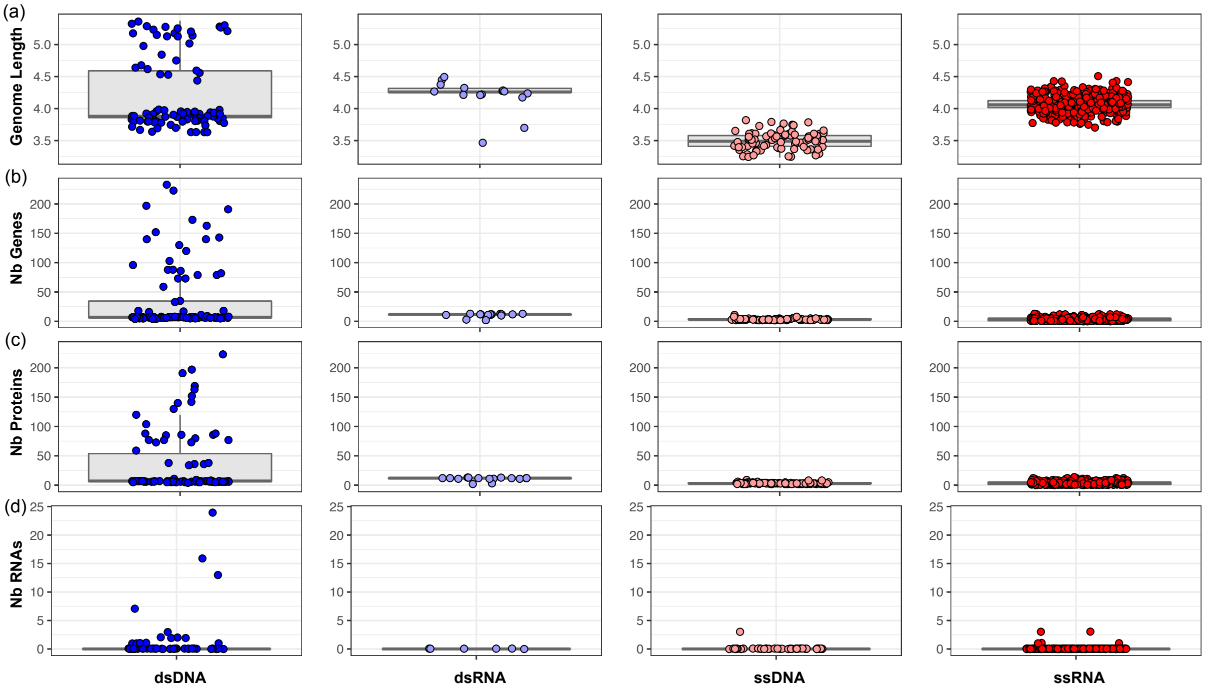


**Figure S2: Physical characteristics of the four types of viruses.** Boxplots show the distribution of four features: (a) genome size, on a $\log_{10}$ scale; (b) number of genes; (c) number of proteins; (d) number of RNAs. These distributions are shown for dsDNA (blue), dsRNA (purple), ssDNA (orange), and ssRNA (red).

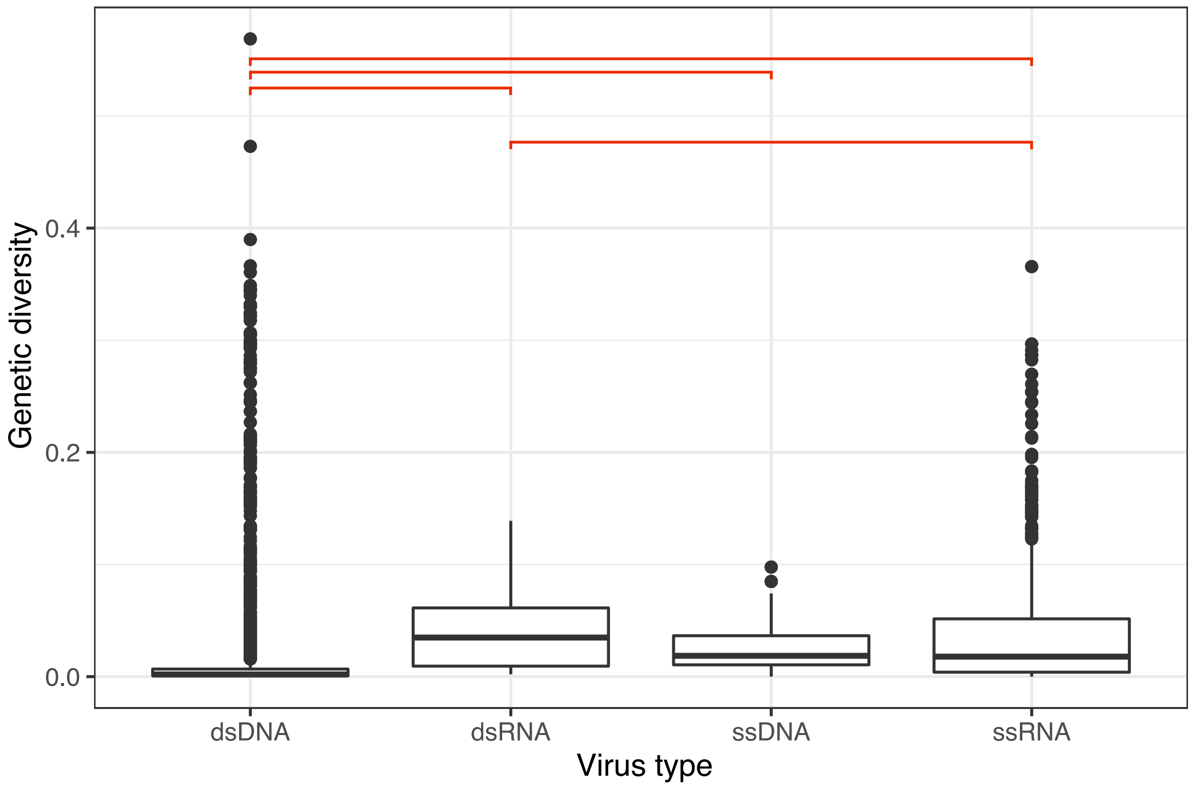


**Figure S3: Viral diversity across viral types.** Boxplots show the distribution of genetic diversity for each viral type, where diversity of each virus of each type is measured by computing the total tree length, divided by the number of sequences in each alignment. Similar results were obtained with mean patristic distances divided by number of sequences. Red bar indicate significant differences at the 1% false discovery rate level (Dunn test; Benjamini-Hochberg correction).

**Table S1: GO enrichment tests for all the MEME genes of interests.** Both tests are based on Fisher's exact test, either in its standard form ("classicFisher"), or in a version designed to be more conservative ("elimFisher"). $P < 0.01$ in bold. Broad viral stages (GO.ID column): E: cell entry/exit; R: replication; T: transcription; A: assembly.

| VirTyp | GO | GO.ID | Term | Annotated | Significant | Expected | Rank in classicFisher | classicFisher | elimFisher |
|---|---|---|---|---|---|---|---|---|---|
| dsDNA | MF | **GO:0004003 (R)** | **ATP-dependent DNA helicase activity** | **122** | **70** | **20.94** | **1** | $\mathbf{1.80\times10^{-26}}$ | $\mathbf{1.80\times10^{-26}}$ |
| | | **GO:0003677 (R)** | **DNA binding** | **343** | **107** | **58.86** | **9** | $\mathbf{4.30\times10^{-14}}$ | $\mathbf{4.30\times10^{-14}}$ |
| | | **GO:0003724 (R)** | **RNA helicase activity** | **149** | **60** | **25.57** | **14** | $\mathbf{8.30\times10^{-13}}$ | $\mathbf{8.30\times10^{-13}}$ |
| | | **GO:0005524** | **ATP binding** | **353** | **94** | **60.58** | **20** | $\mathbf{1.10\times10^{-7}}$ | $\mathbf{1.10\times10^{-7}}$ |
| | | **GO:0003723 (R)** | **RNA binding** | **262** | **64** | **44.96** | **32** | **0.00056** | **0.00056** |
| | | **GO:0003700 (T)** | **DNA binding transcription factor activit...** | **64** | **20** | **10.98** | **35** | **0.00338** | **0.00338** |
| | | **GO:0019783 (R)** | **ubiquitin-like protein-specific protease...** | **5** | **4** | **0.86** | **36** | **0.00367** | **0.00367** |
| | | **GO:0005515 (E)** | **protein binding** | **223** | **52** | **38.27** | **37** | **0.00635** | **0.00635** |
| | | GO:0016773 | phosphotransferase activity alcohol gro | 32 | 11 | 5.49 | 39 | 0.01332 | 0.01332 |
| | | GO:0005506 | iron ion binding | 10 | 5 | 1.72 | 40 | 0.01711 | 0.01711 |
| | BP | **GO:0006260 (R)** | **DNA replication** | **184** | **84** | **24.83** | **1** | $\mathbf{<1\times10^{-30}}$ | $\mathbf{5.30\times10^{-30}}$ |
| | | **GO:0019079 (R)** | **viral genome replication** | **216** | **71** | **29.15** | **3** | $\mathbf{1.00\times10^{-16}}$ | $\mathbf{1.00\times10^{-16}}$ |
| | | **GO:0006275 (R)** | **regulation of DNA replication** | **25** | **11** | **3.37** | **28** | **0.00016** | **0.00016** |
| | | **GO:0022900** | **electron transport chain** | **7** | **5** | **0.94** | **31** | **0.00071** | **0.00071** |
| | | **GO:0040011 (E)** | **locomotion** | **14** | **7** | **1.89** | **32** | **0.00108** | **0.00108** |
| | | **GO:1901566 (R)** | **organonitrogen compound biosynthetic pro...** | **37** | **11** | **4.99** | **34** | **0.00698** | **0.00698** |
| | | **GO:0044283 (R)** | **small molecule biosynthetic process** | **7** | **4** | **0.94** | **35** | **0.00806** | **0.00806** |
| | | **GO:0006383 (T)** | **transcription from RNA polymerase III pr...** | **4** | **3** | **0.54** | **36** | **0.00871** | **0.00871** |
| | | GO:0002252 | immune effector process | 2 | 2 | 0.27 | 38 | 0.01811 | 0.01811 |
| | | GO:0009072 | aromatic amino acid family metabolic pro... | 2 | 2 | 0.27 | 39 | 0.01811 | 0.01811 |
| | CC | **GO:0042025 (R)** | **host cell nucleus** | **66** | **20** | **6.66** | **1** | $\mathbf{1.50\times10^{-6}}$ | $\mathbf{1.50\times10^{-6}}$ |
| | | **GO:0005666 (T)** | **DNA-directed RNA polymerase III complex** | **4** | **3** | **0.4** | **16** | **0.0037** | **0.0037** |
| | | **GO:0005882 (R)** | **intermediate filament** | **4** | **3** | **0.4** | **17** | **0.0037** | **0.0037** |
| | | GO:0033179 | proton-transporting V-type ATPase V0 do | 2 | 2 | 0.2 | 23 | 0.0101 | 0.0101 |
| | | GO:0045095 | keratin filament | 2 | 2 | 0.2 | 24 | 0.0101 | 0.0101 |
| | | GO:0046806 | viral scaffold | 2 | 2 | 0.2 | 25 | 0.0101 | 0.0101 |
| | | GO:0042597 | periplasmic space | 12 | 4 | 1.21 | 29 | 0.0255 | 0.0255 |
| | | GO:0005887 | integral component of plasma membrane | 3 | 2 | 0.3 | 31 | 0.0282 | 0.0282 |
| | | GO:0005929 | cilium | 3 | 2 | 0.3 | 32 | 0.0282 | 0.0282 |
| | | GO:0031226 | intrinsic component of plasma membrane | 3 | 2 | 0.3 | 33 | 0.0282 | 0.0282 |
| ssRNA | MF | **GO:0003968 (R)** | **RNA-directed 5'-3' RNA polymerase activi...** | **257** | **84** | **28.5** | **1** | $\mathbf{3.90\times10^{-27}}$ | $\mathbf{3.90\times10^{-27}}$ |
| | | **GO:0004197 (R)** | **cysteine-type endopeptidase activity** | **52** | **28** | **5.77** | **10** | $\mathbf{8.00\times10^{-15}}$ | $\mathbf{8.00\times10^{-15}}$ |
| | | **GO:0004252 (R)** | **serine-type endopeptidase activity** | **79** | **29** | **8.76** | **18** | $\mathbf{4.90\times10^{-10}}$ | $\mathbf{4.90\times10^{-10}}$ |
| | | **GO:0046983 (R)** | **protein dimerization activity** | **38** | **18** | **4.21** | **20** | $\mathbf{1.20\times10^{-8}}$ | $\mathbf{1.20\times10^{-8}}$ |
| | | **GO:0046789 (E)** | **host cell surface receptor binding** | **31** | **16** | **3.44** | **21** | $\mathbf{1.70\times10^{-8}}$ | $\mathbf{1.70\times10^{-8}}$ |
| | | **GO:0004482 (R)** | **mRNA (guanine-N7-)-methyltransferase act...** | **86** | **26** | **9.54** | **25** | $\mathbf{4.50\times10^{-7}}$ | $\mathbf{4.50\times10^{-7}}$ |
| | | **GO:0005198 (R)** | **structural molecule activity** | **327** | **62** | **36.26** | **28** | $\mathbf{7.20\times10^{-7}}$ | $\mathbf{7.20\times10^{-7}}$ |
| | | **GO:0003725 (R)** | **double-stranded RNA binding** | **28** | **13** | **3.1** | **30** | $\mathbf{2.00\times10^{-6}}$ | $\mathbf{2.00\times10^{-6}}$ |
| | | **GO:0004483 (R)** | **mRNA (nucleoside-2'-O-)-methyltransferas...** | **27** | **12** | **2.99** | **32** | $\mathbf{9.10\times10^{-6}}$ | $\mathbf{9.10\times10^{-6}}$ |
| | | **GO:0008168 (R)** | **methyltransferase activity** | **113** | **37** | **12.53** | **16** | $\mathbf{6.30\times10^{-11}}$ | $\mathbf{1.90\times10^{-5}}$ |
| | BP | **GO:0006508 (R)** | **proteolysis** | **120** | **49** | **15.71** | **2** | $\mathbf{4.40\times10^{-16}}$ | $\mathbf{4.40\times10^{-16}}$ |
| | | **GO:0032259 (R)** | **methylation** | **66** | **29** | **8.64** | **11** | $\mathbf{1.60\times10^{-10}}$ | $\mathbf{1.60\times10^{-10}}$ |
| | | **GO:0019087 (R)** | **transformation of host cell by virus** | **43** | **21** | **5.63** | **14** | $\mathbf{6.90\times10^{-9}}$ | $\mathbf{6.90\times10^{-9}}$ |
| | | **GO:0019082 (R)** | **viral protein processing** | **20** | **13** | **2.62** | **16** | $\mathbf{7.20\times10^{-8}}$ | $\mathbf{7.20\times10^{-8}}$ |
| | | **GO:0019058** | **viral life cycle** | **411** | **87** | **53.81** | **13** | $\mathbf{6.70\times10^{-9}}$ | $\mathbf{1.20\times10^{-5}}$ |
| | | **GO:0016070** | **RNA metabolic process** | **413** | **76** | **54.07** | **17** | $\mathbf{8.90\times10^{-5}}$ | $\mathbf{8.90\times10^{-5}}$ |
| | | **GO:0019064 (E)** | **fusion of virus membrane with host plasm...** | **48** | **16** | **6.28** | **18** | **0.00019** | **0.00019** |
| | | **GO:0019048** | **modulation by virus of host morphology o...** | **84** | **39** | **11** | **3** | $\mathbf{5.10\times10^{-15}}$ | **0.00021** |
| | | **GO:0030683** | **evasion or tolerance by virus of host im...** | **24** | **10** | **3.14** | **29** | **0.00043** | **0.00043** |
| | | **GO:0039694 (R)** | **viral RNA genome replication** | **13** | **7** | **1.7** | **34** | **0.0005** | **0.0005** |
| | CC | **GO:0019031** | **viral envelope** | **164** | **51** | **25.27** | **2** | $\mathbf{9.60\times10^{-9}}$ | $\mathbf{9.60\times10^{-9}}$ |
| | | **GO:0019012** | **virion** | **601** | **128** | **92.62** | **1** | $\mathbf{3.20\times10^{-14}}$ | $\mathbf{3.00\times10^{-8}}$ |
| | | **GO:0055036** | **virion membrane** | **24** | **11** | **3.7** | **5** | **0.00033** | **0.00033** |
| | | GO:0044454 | nuclear chromosome part | 11 | 5 | 1.7 | 6 | 0.01712 | 0.01712 |
| | | GO:0000778 | condensed nuclear chromosome kinetochore | 10 | 4 | 1.54 | 7 | 0.05346 | 0.05346 |
| | | GO:0000780 | condensed nuclear chromosome centromeri | 10 | 4 | 1.54 | 8 | 0.05346 | 0.05346 |
| | | GO:0000942 | condensed nuclear chromosome outer kinet... | 10 | 4 | 1.54 | 9 | 0.05346 | 0.05346 |
| | | GO:0042729 | DASH complex | 10 | 4 | 1.54 | 10 | 0.05346 | 0.05346 |
| | | GO:0072686 | mitotic spindle | 10 | 4 | 1.54 | 11 | 0.05346 | 0.05346 |
| | | GO:0019030 | icosahedral viral capsid | 3 | 2 | 0.46 | 12 | 0.06362 | 0.06362 |

**Table S2: GO enrichment tests for all the SM genes of interests.** Both tests are based on Fisher's exact test, either in its standard form ("classicFisher"), or in a version designed to be more conservative ("elimFisher"). $P < 0.01$ in bold. Broad viral stages (GO.ID column): E: cell entry/exit; R: replication; T: transcription; A: assembly.

| VirTyp | GO | GO.ID | Term | Annotated | Significant | Expected | Rank in classicFisher | classicFisher | elimFisher |
|---|---|---|---|---|---|---|---|---|---|
| dsDNA | MF | **GO:0004003 (R)** | **ATP-dependent DNA helicase activity** | **122** | **51** | **28.17** | **1** | $\mathbf{1.10\times10^{-6}}$ | $\mathbf{1.10\times10^{-6}}$ |
| | | **GO:0003677 (R)** | **DNA binding** | **343** | **109** | **79.21** | **4** | $\mathbf{1.30\times10^{-5}}$ | $\mathbf{1.30\times10^{-5}}$ |
| | | **GO:0050662 (R)** | **coenzyme binding** | **10** | **8** | **2.31** | **8** | **0.00022** | **0.00022** |
| | | **GO:0016616 (E)** | **oxidoreductase activity acting on the C** | **4** | **4** | **0.92** | **9** | **0.0028** | **0.0028** |
| | | **GO:0016984** | **ribulose-bisphosphate carboxylase activi...** | **6** | **5** | **1.39** | **10** | **0.00312** | **0.00312** |
| | | **GO:0004484 (T)** | **mRNA guanylyltransferase activity** | **11** | **7** | **2.54** | **14** | **0.00456** | **0.00456** |
| | | **GO:0020037 (E)** | **heme binding** | **11** | **7** | **2.54** | **15** | **0.00456** | **0.00456** |
| | | **GO:0016773 (R)** | **phosphotransferase activity alcohol gro** | **32** | **14** | **7.39** | **21** | **0.00715** | **0.00715** |
| | | **GO:0031423** | **hexon binding** | **7** | **5** | **1.62** | **23** | **0.00885** | **0.00885** |
| | | GO:0019783 | ubiquitin-like protein-specific protease... | 5 | 4 | 1.15 | 24 | 0.01145 | 0.01145 |
| | BP | **GO:0006260 (R)** | **DNA replication** | **184** | **59** | **40.49** | **2** | **0.00039** | **0.00039** |
| | | **GO:0015977 (R)** | **carbon fixation** | **6** | **5** | **1.32** | **3** | **0.00247** | **0.00247** |
| | | **GO:0022900** | **electron transport chain** | **7** | **5** | **1.54** | **5** | **0.00708** | **0.00708** |
| | | **GO:0044283 (R)** | **small molecule biosynthetic process** | **7** | **5** | **1.54** | **6** | **0.00708** | **0.00708** |
| | | **GO:1901566 (R)** | **organonitrogen compound biosynthetic pro...** | **37** | **15** | **8.14** | **7** | **0.00771** | **0.00771** |
| | | **GO:0007166** | **cell surface receptor signaling pathway** | **5** | **4** | **1.1** | **9** | **0.00952** | **0.00952** |
| | | GO:0007276 | gamete generation | 10 | 6 | 2.2 | 10 | 0.01012 | 0.01012 |
| | | GO:0019953 | sexual reproduction | 10 | 6 | 2.2 | 11 | 0.01012 | 0.01012 |
| | | GO:0032504 | multicellular organism reproduction | 10 | 6 | 2.2 | 12 | 0.01012 | 0.01012 |
| | | GO:0044703 | multi-organism reproductive process | 10 | 6 | 2.2 | 13 | 0.01012 | 0.01012 |
| | CC | **GO:0042597 (E)** | **periplasmic space** | **12** | **7** | **1.98** | **5** | **0.00112** | **0.0011** |
| | | **GO:0043229** | **intracellular organelle** | **101** | **27** | **16.68** | **7** | **0.00389** | **0.0039** |
| | | **GO:0032991** | **macromolecular complex** | **96** | **28** | **15.86** | **3** | **0.00072** | **0.004** |
| | | **GO:0030089** | **phycobilisome** | **3** | **3** | **0.5** | **8** | **0.00443** | **0.0044** |
| | | **GO:0031514 (E)** | **motile cilium** | **3** | **3** | **0.5** | **9** | **0.00443** | **0.0044** |
| | | GO:0019033 | viral tegument | 10 | 5 | 1.65 | 16 | 0.01436 | 0.0144 |
| | | GO:0000428 | DNA-directed RNA polymerase complex | 4 | 3 | 0.66 | 17 | 0.01557 | 0.0156 |
| | | GO:0005666 | DNA-directed RNA polymerase III complex | 4 | 3 | 0.66 | 18 | 0.01557 | 0.0156 |
| | | GO:0030880 | RNA polymerase complex | 4 | 3 | 0.66 | 19 | 0.01557 | 0.0156 |
| | | GO:0055029 | nuclear DNA-directed RNA polymerase comp... | 4 | 3 | 0.66 | 20 | 0.01557 | 0.0156 |
| ssRNA | MF | **GO:0046789 (E)** | **host cell surface receptor binding** | **31** | **7** | **1.07** | **1** | $\mathbf{5.40\times10^{-5}}$ | $\mathbf{5.40\times10^{-5}}$ |
| | | **GO:0003950 (T)** | **NAD+ ADP-ribosyltransferase activity** | **5** | **3** | **0.17** | **3** | **0.00037** | **0.00037** |
| | | **GO:0005198 (R)** | **structural molecule activity** | **327** | **20** | **11.32** | **7** | **0.00369** | **0.00369** |
| | | **GO:0016798 (E)** | **hydrolase activity acting on glycosyl b** | **12** | **3** | **0.42** | **8** | **0.00688** | **0.00688** |
| | | **GO:0004482 (T)** | **mRNA (guanine-N7-)-methyltransferase act...** | **86** | **8** | **2.98** | **9** | **0.00772** | **0.00772** |
| | | GO:0004308 | exo-alpha-sialidase activity | 6 | 2 | 0.21 | 13 | 0.0161 | 0.0161 |
| | | GO:0016997 | alpha-sialidase activity | 6 | 2 | 0.21 | 14 | 0.0161 | 0.0161 |
| | | GO:0003968 | RNA-directed 5'-3' RNA polymerase activi... | 257 | 15 | 8.89 | 16 | 0.02182 | 0.02182 |
| | | GO:0004553 | hydrolase activity hydrolyzing O-glycos | 10 | 2 | 0.35 | 19 | 0.04422 | 0.04422 |
| | | GO:0000702 | oxidized base lesion DNA N-glycosylase a... | 2 | 1 | 0.07 | 20 | 0.06804 | 0.06804 |
| | BP | **GO:0006471 (T)** | **protein ADP-ribosylation** | **5** | **3** | **0.18** | **1** | **0.00039** | **0.00039** |
| | | **GO:0006352 (T)** | **DNA-templated transcription initiation** | **4** | **2** | **0.14** | **8** | **0.007** | **0.007** |
| | | **GO:0055085 (E)** | **transmembrane transport** | **23** | **4** | **0.81** | **9** | **0.00736** | **0.00736** |
| | | GO:0006370 | 7-methylguanosine mRNA capping | 63 | 6 | 2.23 | 11 | 0.02051 | 0.02051 |
| | | GO:0009452 | 7-methylguanosine RNA capping | 63 | 6 | 2.23 | 12 | 0.02051 | 0.02051 |
| | | GO:0036260 | RNA capping | 63 | 6 | 2.23 | 13 | 0.02051 | 0.02051 |
| | | GO:0019068 | virion assembly | 66 | 6 | 2.33 | 14 | 0.02532 | 0.02532 |
| | | GO:0006397 | mRNA processing | 68 | 6 | 2.4 | 15 | 0.02891 | 0.02891 |
| | | GO:0016071 | mRNA metabolic process | 79 | 6 | 2.79 | 16 | 0.0548 | 0.0548 |
| | | GO:0005975 | carbohydrate metabolic process | 12 | 2 | 0.42 | 17 | 0.06435 | 0.06435 |
| | CC | **GO:0019031 (E)** | **viral envelope** | **164** | **19** | **9.82** | **3** | **0.0016** | **0.0016** |
| | | **GO:0019013** | **viral nucleocapsid** | **104** | **14** | **6.23** | **5** | **0.0019** | **0.0019** |
| | | GO:0019012 | virion | 601 | 49 | 35.98 | 1 | $2.40\times10^{-5}$ | 0.015 |
| | | GO:0055036 | virion membrane | 24 | 4 | 1.44 | 6 | 0.0498 | 0.0498 |
| | | GO:0009654 | photosystem II oxygen evolving complex | 1 | 1 | 0.06 | 7 | 0.0599 | 0.0599 |
| | | GO:0019898 | extrinsic component of membrane | 1 | 1 | 0.06 | 8 | 0.0599 | 0.0599 |
| | | GO:1990204 | oxidoreductase complex | 1 | 1 | 0.06 | 9 | 0.0599 | 0.0599 |
| | | GO:0033180 | proton-transporting V-type ATPase V1 do | 2 | 1 | 0.12 | 10 | 0.1162 | 0.1162 |
| | | GO:0033644 | host cell membrane | 12 | 2 | 0.72 | 11 | 0.1582 | 0.1582 |
| | | GO:0009521 | photosystem | 3 | 1 | 0.18 | 12 | 0.1692 | 0.1692 |

**Table S3: GO enrichment tests for all the SM+MEME genes of interests.** Both tests are based on Fisher's exact test, either in its standard form ("classicFisher"), or in a version designed to be more conservative ("elimFisher"). $P < 0.01$ in bold. Broad viral stages (GO.ID column): E: cell entry/exit; R: replication; T: transcription; A: assembly.

| VirTyp | GO | GO.ID | Term | Annotated | Significant | Expected | Rank in classicFisher | classicFisher | elimFisher |
|---|---|---|---|---|---|---|---|---|---|
| dsDNA | MF | **GO:0005198 (R)** | **structural molecule activity** | **327** | **9** | **2.54** | **1** | **$6.00\times10^{-5}$** | **$6.00\times10^{-5}$** |
| | | GO:0004252 | serine-type endopeptidase activity | 79 | 2 | 0.61 | 2 | 0.12 | 0.12 |
| | | GO:0008236 | serine-type peptidase activity | 90 | 2 | 0.7 | 3 | 0.15 | 0.15 |
| | | GO:0017171 | serine hydrolase activity | 90 | 2 | 0.7 | 4 | 0.15 | 0.15 |
| | | GO:0004175 | endopeptidase activity | 133 | 2 | 1.03 | 5 | 0.28 | 0.28 |
| | | GO:0070011 | peptidase activity acting on L-amino ac | 149 | 2 | 1.16 | 6 | 0.32 | 0.32 |
| | | GO:0008233 | peptidase activity | 153 | 2 | 1.19 | 7 | 0.34 | 0.34 |
| | | GO:0140096 | catalytic activity acting on a protein | 191 | 2 | 1.48 | 8 | 0.45 | 0.45 |
| | | GO:0003677 | DNA binding | 343 | 2 | 2.66 | 9 | 0.79 | 0.79 |
| | | GO:0016787 | hydrolase activity | 402 | 2 | 3.12 | 10 | 0.86 | 0.86 |
| | BP | **GO:0043158** | **heterocyst differentiation** | **6** | **2** | **0.02** | **1** | **0.00012** | **0.00012** |
| | | **GO:0051726** | **regulation of cell cycle** | **7** | **2** | **0.02** | **2** | **0.00016** | **0.00016** |
| | | GO:0007049 | cell cycle | 29 | 2 | 0.09 | 6 | 0.00306 | 1 |
| | | GO:0008150 | biological_process | 1245 | 4 | 4 | 11 | 1 | 1 |
| | | GO:0009987 | cellular process | 809 | 4 | 2.6 | 10 | 0.17782 | 1 |
| | | GO:0030154 | cell differentiation | 15 | 2 | 0.05 | 3 | 0.0008 | 1 |
| | | GO:0032502 | developmental process | 20 | 2 | 0.06 | 5 | 0.00144 | 1 |
| | | GO:0048869 | cellular developmental process | 15 | 2 | 0.05 | 4 | 0.0008 | 1 |
| | | GO:0050789 | regulation of biological process | 225 | 2 | 0.72 | 8 | 0.15168 | 1 |
| | | GO:0050794 | regulation of cellular process | 206 | 2 | 0.66 | 7 | 0.12999 | 1 |
| | CC | **GO:0019028 (E)** | **viral capsid** | **375** | **9** | **4.57** | **1** | **0.0077** | **0.0077** |
| | | GO:0005634 | nucleus | 49 | 2 | 0.6 | 3 | 0.1163 | 0.1163 |
| | | GO:0043231 | intracellular membrane-bounded organelle | 72 | 2 | 0.88 | 4 | 0.2169 | 0.2169 |
| | | GO:0043227 | membrane-bounded organelle | 80 | 2 | 0.98 | 6 | 0.2543 | 0.2543 |
| | | GO:0043229 | intracellular organelle | 101 | 2 | 1.23 | 7 | 0.3538 | 0.3538 |
| | | GO:0043226 | organelle | 110 | 2 | 1.34 | 8 | 0.3959 | 0.3959 |
| | | GO:0044424 | intracellular part | 136 | 2 | 1.66 | 9 | 0.5119 | 0.5119 |
| | | GO:0005622 | intracellular | 138 | 2 | 1.68 | 10 | 0.5203 | 0.5203 |
| | | GO:0005623 | cell | 171 | 2 | 2.09 | 11 | 0.6477 | 0.6477 |
| | | GO:0044464 | cell part | 171 | 2 | 2.09 | 12 | 0.6477 | 0.6477 |
| ssRNA | MF | **GO:0046789 (E)** | **host cell surface receptor binding** | **31** | **6** | **0.46** | **1** | **$2.90\times10^{-6}$** | **$2.90\times10^{-6}$** |
| | | GO:0005509 | calcium ion binding | 2 | 1 | 0.03 | 5 | 0.029 | 0.029 |
| | | GO:0005198 | structural molecule activity | 327 | 9 | 4.85 | 6 | 0.034 | 0.034 |
| | | GO:0005200 | structural constituent of cytoskeleton | 3 | 1 | 0.04 | 7 | 0.044 | 0.044 |
| | | GO:0004483 | mRNA (nucleoside-2'-O-)-methyltransferas... | 27 | 2 | 0.4 | 8 | 0.059 | 0.059 |
| | | GO:0008171 | O-methyltransferase activity | 27 | 2 | 0.4 | 9 | 0.059 | 0.059 |
| | | GO:0070008 | serine-type exopeptidase activity | 27 | 2 | 0.4 | 10 | 0.059 | 0.059 |
| | | GO:0003725 | double-stranded RNA binding | 28 | 2 | 0.42 | 11 | 0.063 | 0.063 |
| | | GO:0003950 | NAD+ ADP-ribosyltransferase activity | 5 | 1 | 0.07 | 12 | 0.072 | 0.072 |
| | | GO:0016763 | transferase activity transferring pento | 5 | 1 | 0.07 | 13 | 0.072 | 0.072 |
| | BP | **GO:0006352 (T)** | **DNA-templated transcription initiation** | **4** | **2** | **0.06** | **1** | **0.0013** | **0.0013** |
| | | **GO:0019064 (E)** | **fusion of virus membrane with host plasm...** | **48** | **4** | **0.73** | **3** | **0.0049** | **0.0049** |
| | | GO:0007155 | cell adhesion | 16 | 2 | 0.24 | 16 | 0.0233 | 0.0233 |
| | | GO:0006891 | intra-Golgi vesicle-mediated transport | 3 | 1 | 0.05 | 17 | 0.0451 | 0.0451 |
| | | GO:0019058 | viral life cycle | 411 | 12 | 6.27 | 13 | 0.0064 | 0.052 |
| | | GO:0022610 | biological adhesion | 27 | 2 | 0.41 | 22 | 0.0617 | 0.0617 |
| | | GO:0006471 | protein ADP-ribosylation | 5 | 1 | 0.08 | 24 | 0.0741 | 0.0741 |
| | | GO:0006486 | protein glycosylation | 5 | 1 | 0.08 | 25 | 0.0741 | 0.0741 |
| | | GO:0009100 | glycoprotein metabolic process | 5 | 1 | 0.08 | 26 | 0.0741 | 0.0741 |
| | | GO:0009101 | glycoprotein biosynthetic process | 5 | 1 | 0.08 | 27 | 0.0741 | 0.0741 |
| | CC | **GO:0019031 (E)** | **viral envelope** | **164** | **12** | **4.36** | **1** | **0.00032** | **0.00032** |
| | | GO:0009654 | photosystem II oxygen evolving complex | 1 | 1 | 0.03 | 3 | 0.02661 | 0.02661 |
| | | GO:0019898 | extrinsic component of membrane | 1 | 1 | 0.03 | 4 | 0.02661 | 0.02661 |
| | | GO:1990204 | oxidoreductase complex | 1 | 1 | 0.03 | 5 | 0.02661 | 0.02661 |
| | | GO:0043234 | protein complex | 72 | 5 | 1.92 | 6 | 0.03611 | 0.03611 |
| | | GO:0033180 | proton-transporting V-type ATPase V1 do | 2 | 1 | 0.05 | 7 | 0.05254 | 0.05254 |
| | | GO:0009521 | photosystem | 3 | 1 | 0.08 | 8 | 0.0778 | 0.0778 |
| | | GO:0009523 | photosystem II | 3 | 1 | 0.08 | 9 | 0.0778 | 0.0778 |
| | | GO:0017119 | Golgi transport complex | 3 | 1 | 0.08 | 10 | 0.0778 | 0.0778 |
| | | GO:0044431 | Golgi apparatus part | 3 | 1 | 0.08 | 11 | 0.0778 | 0.0778 |
